# TRUHiC: A TRansformer-embedded U-2 Net to enhance Hi-C data for 3D chromatin structure characterization

**DOI:** 10.1101/2025.03.29.646133

**Authors:** Chong Li, Mohammad Erfan Mowlaei, Lingbin Ni, Human Genome Structural Variation Consortium (HGSVC), HGSVC Functional Analysis Working Group, Mark J.P. Chaisson, Evan E. Eichler, Vincenzo Carnevale, Sudhir Kumar, Xinghua Shi

**Affiliations:** Department of Computer and Information Sciences, College of Science and Technology, Temple University, Philadelphia, PA, US; Institute for Genomics and Evolutionary Medicine, Temple University 1925 N. 12th Street, 19122, PA, USA; Institute for Computational Molecular Science, Temple University 1925 N. 12th Street, 19122, PA, USA; Department of Biology, College of Science and Technology, Temple University, Philadelphia, PA, USA; Department of Quantitative and Computational Biology, University of Southern California, Los Angeles, CA, USA; Department of Genome Sciences, University of Washington School of Medicine, Seattle, WA, USA; Howard Hughes Medical Institute, University of Washington, Seattle, WA, USA

**Keywords:** Hi-C, 3D genome, deep learning, transformer, super-resolution

## Abstract

High-throughput chromosome conformation capture sequencing (Hi-C) is a key technology for studying the three-dimensional (3D) structure of genomes and chromatin folding. Hi-C data reveals underlying patterns of genome organization, such as topologically associating domains (TADs) and chromatin loops, with critical roles in transcriptional regulation and disease etiology and progression. However, the sparsity of existing Hi-C data often hinders robust and reliable inference of 3D structures. Hence, we propose *TRUHiC*, a new computational method that leverages recent state-of-the-art deep generative modeling to augment low-resolution Hi-C data for the characterization of 3D chromatin structures. By applying *TRUHiC* to real low-resolution Hi-C data from the GM12329 cell line and across other publicly available Hi-C data for human and mice, we demonstrate that the augmented data significantly improve the characterization of TADs and loops across diverse cell lines and species. We further present a pre-trained *TRUHiC* on human lymphoblastoid cell lines that can be adaptable and transferable to improve chromatin characterization of various cell lines, tissues, and species.

## 1 Introduction

The technology of high-throughput chromosome conformation capture sequencing (Hi-C) has emerged as a pivotal approach for studying the three-dimensional (3D) genome organization^1^. This technology builds upon the principles of chromatin conformation capture assay (3C) and enables researchers to explore the interactions of chromatin across the entire genome. The analysis of Hi-C data at desired resolutions would facilitate a comprehensive understanding of genome-wide chromatin structures, such as A/B compartments^1^, topologically associating domains (TADs)^2^, and chromatin loops^3^, thereby shedding light on the essential functions of the 3D genome^2^. Nevertheless, biologically meaningful identification of fine-grained structural features, especially TADs and loops, necessitates higher resolution or read depth of Hi-C sequencing. Achieving this resolution demands costly high-coverage deep sequencing to ensure sufficient read depth for accurately capturing chromatin interaction frequencies^4^. Consequently, many existing Hi-C datasets present relatively low resolution, defined by larger genomic bins (fixed-length windows, e.g., 10 kb, 50 kb, 100 kb) partitioned across the genome due to cost constraints limiting their utility in discerning finer chromatin structures^5,6^. This issue has prompted the development of computational methods for Hi-C data enhancement to leverage existing low-resolution Hi-C data to infer corresponding high-resolution Hi-C data for various genomic downstream analyses, such as identifying enhancer–promoter interactions or mapping topologically associating domains, thereby enabling detailed investigations into how the spatial organization of chromatin influences gene regulatory mechanisms by modulating the accessibility of DNA to transcriptional machinery^7^.

The computational enhancement of Hi-C data shares conceptual similarities with the image super-resolution task in computer vision^8,9^. Hi-C sequencing reads are typically transformed into contact matrices that can be visualized as image-like heatmaps, suggesting a potential application of super-resolution techniques (Figure 1a). However, directly applying image-based super-resolution methods to Hi-C contact maps is less effective due to the unique structural properties of Hi-C contact matrices. Unlike natural images, Hi-C contact matrices are inherently symmetric around the diagonal and contain biologically meaningful constraints, such as sparsity in long-range interactions and high variability across genomic regions. The number of rows and columns in a Hi-C matrix corresponds to the length of the genome divided by the resolution (bin size, smaller bin size of fixed genomic intervals)^10^. Key 3D chromatin features such as TADs are visually represented as triangular regions with elevated signal intensity on the heat maps, and chromatin loops are depicted as concentrated focal points (Figure 1a)^11^. Thus, developing specialized Hi-C resolution enhancement methods that incorporate biological domain knowledge is essential for accurately reconstructing high-resolution Hi-C contact maps.

**Figure 1.**
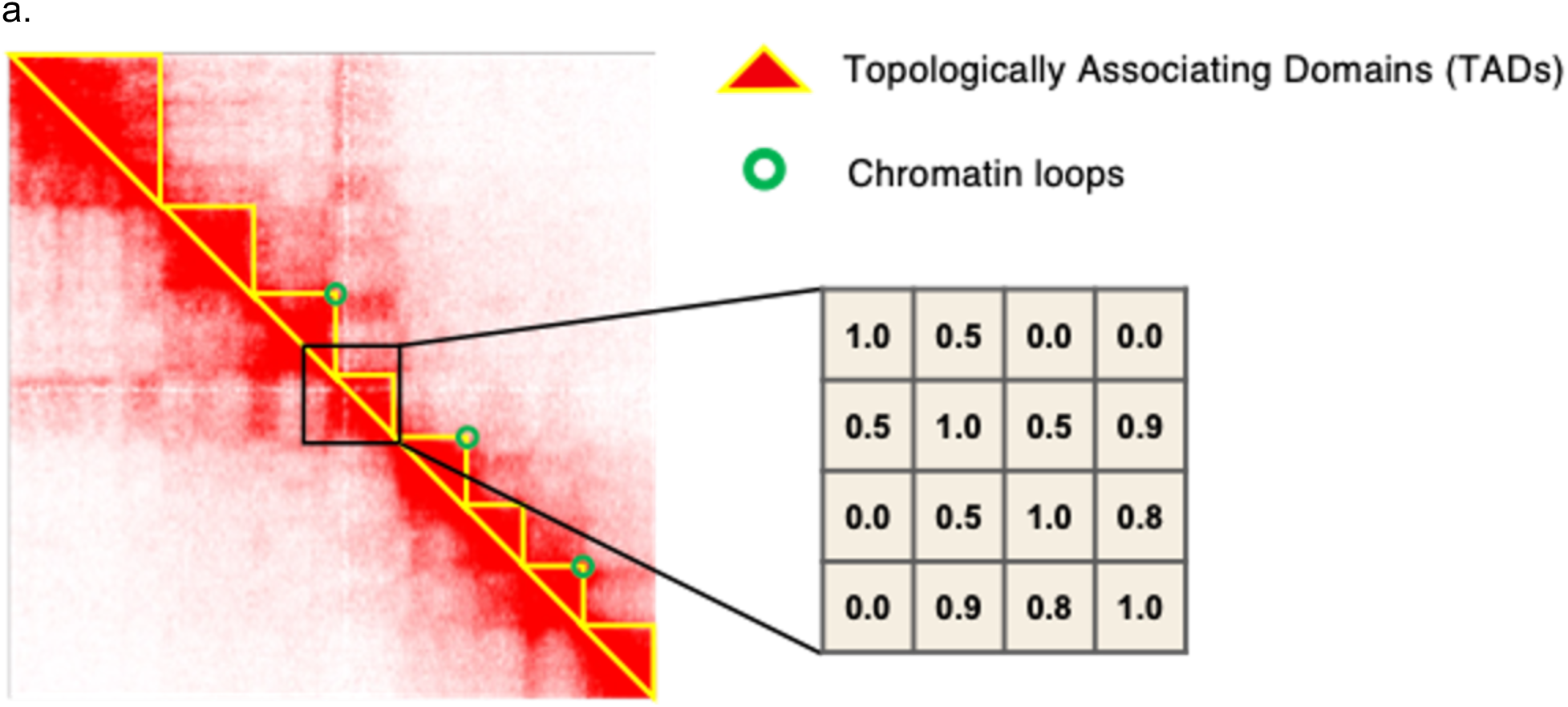
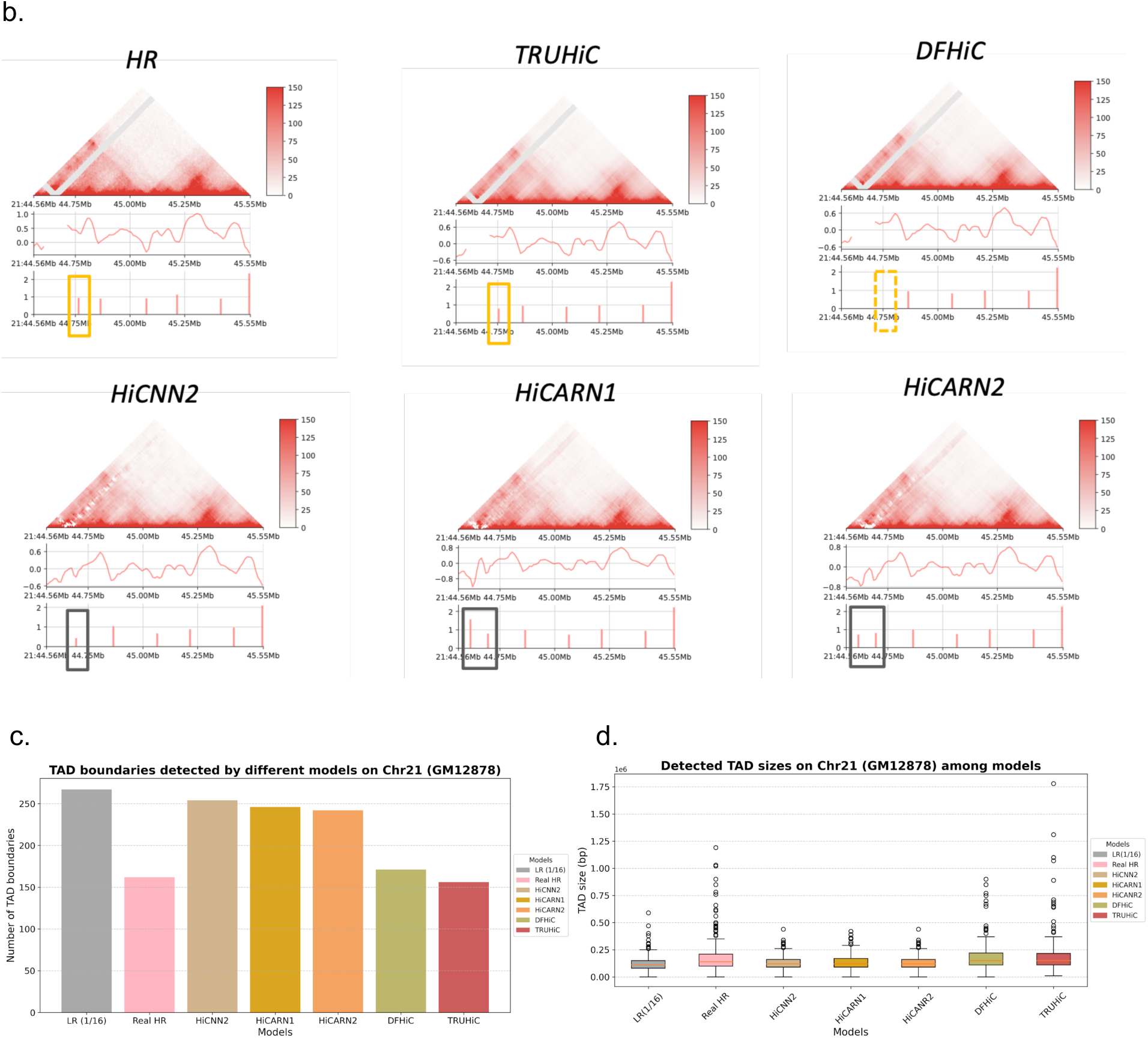
The performance comparison of the *TRUHiC* and its competing methods. **a).** A Hi-C contact map visualizing chromatin interactions, with TAD boundaries outlined in yellow and chromatin loops marked with green circles. The hierarchical organization of chromatin interactions is depicted, highlighting the structural features that contribute to genome organization. A zoomed-in region of the Hi-C contact map displays a representative interaction frequency (IF) matrix, where each value quantifies the frequency of interactions between genomic loci. Higher IF values indicate stronger chromatin interactions. **b).** Visualization of the result in the genomic region Chr 21: 44,560,001-45,550,000, where *TRUHiC* identified the most consistent TAD boundary locations in the real high-resolution data, compared to the competing methods. Yellow rectangles highlight TAD boundary locations detected in both the HR Hi-C contact map and the enhanced maps, while dashed yellow rectangles indicate TAD boundary locations present in the HR Hi-C contact map but missing in the enhanced maps. Black rectangles indicate erroneously detected TAD boundaries that are absent in the HR Hi-C contact map but appear in the enhanced maps. **c-d).** Comparison of the number of TAD boundaries and sizes of TADs detected using *IS* on chromosome 21 of the GM12878 cell line recovered by different methods for down-sampled low-resolution Hi-C data. The results for other test chromosomes are presented in Supplementary Figures S1 and S2.

Various deep learning methods have been developed to address the resolution enhancement challenge, predominantly based on two architectures: convolutional neural networks (CNNs) and generative adversarial networks (GANs), as well as their variants. Existing CNN-based methods include *HiCPlus*^5^, *HiCNN*^8^ and *HiCNN2*^9^, *SRHiC*^12^, and *DFHiC*^13^, while current GAN-based methods contain *hicGAN*^6^, *DeepHiC*^14^, *HiCSR*^15^, *VEHiCLE*^16^, and *EnHiC*^17^, accompanied by an integrated CNN and GAN-based model termed *HiCARN*^18^. These methods differ in terms of the loss function and model architecture that led to varying performances. *HiCPlus* pioneered the use of CNN architecture with mean squared error (MSE) loss, while *HiCNN* introduced a deeper convolutional network. *hicGAN* was the first to incorporate a GAN framework to generate high-resolution contact maps conditioned on low-resolution inputs. *HiCNN2* proposed three customized CNN architectures, and *SRHiC* later combined ResNet^19^ and *WDSR* architectures^20^. Subsequent models, such as *DeepHiC* and *HiCSR,* introduced multiple loss components to improve performance. *VEHiCLE* employed a variational autoencoder within a conditional GAN framework, while *EnHiC*^21^ addressed the issue of image artifacts through a novel decomposition and reconstruction block. *HiCARN* employed a lightweight Cascading Residual Network (CARN)^22^, and the latest method, *DFHiC,* integrated peripheral genomic information through their implementation of convolution layers. Beyond these, genome graph-based approaches have recently emerged to infer genome sequences from Hi-C reads and generate more accurate Hi-C contact matrices^23^.

The aforementioned methods primarily adopt an image super-resolution approach, where the Hi-C matrix is segmented, independently processed for resolution enhancement, and then reassembled to construct the enhanced full contact map. However, this approach does not fully account for the biological significance of Hi-C contact matrices, where interaction intensities are inherently tied to the hierarchical nature of 3D chromatin organization. Existing methods rely heavily on convolutional layers. Convolution layers can effectively model local patterns but fall short in capturing long-range chromatin interactions and global structural context. As a result, enhanced contact maps often lack structural coherence and biological fidelity. These limitations underscore the need for new approaches capable of integrating both local and global dependencies. Transformer-based approaches have achieved major breakthroughs in computer vision and natural language processing. However, the quadratic computational cost of the attention mechanism makes it prohibitive to apply it directly to image-like inputs with thousands of features. A solution is to use CNNs to capture local patterns and reduce the dimensionality of the input before using transformer blocks^24^.

Based on these principles, we propose a novel Hi-C data resolution enhancement approach (*TRUHiC*) that integrates a customized and lightweight CNN-based U-2 Net architecture empowered by a transformer block^25,26^. This hybrid architecture leverages the strengths of both convolutional and attention-based mechanisms. The U-2 Net’s deeply nested structure is well-suited for capturing fine-grained, multi-scale local features in contact maps while maintaining a low computational overhead, an important consideration for high-throughput genomic data. At the same time, the transformer block enables the model to effectively capture long-range dependencies and complex interaction patterns, which are essential for reconstructing 3D genome organization. To the best of our knowledge, this is one of the first methods to harness the transformer’s attention mechanism for capturing global chromatin interaction patterns from low-resolution Hi-C contact matrices. We demonstrate that our method outperforms state-of-the-art techniques in both contact map generation and chromatin structure identification, as evidenced by superior evaluation scores across various experimental settings. Understanding these interactions at a finer scale is crucial for improving 3D genome reconstruction and enabling more accurate downstream biological interpretations.

Beyond architectural innovation, a key application of *TRUHiC* is its ability to impute and substantially improve real low-coverage data. For example, when applied to the low-coverage Hi-C dataset of GM12329, *TRUHiC* markedly enhanced TAD boundary detection and loop identification, recovering structural features that are almost entirely missed at the original sequencing depth. This demonstrates its immediate practical value for extending analyses to samples where deep sequencing is either technically infeasible or cost-prohibitive.

Furthermore, existing models, which are typically trained on artificially low-resolution datasets, often experience a dramatic drop in performance when applied to biological replications. To overcome this limitation^27^, we propose *TRUHiC-LCL, a* cell-line-specific *(CLS)* model for lymphoblastoid cell lines (LCLs) trained on a large real LCL-specific Hi-C dataset (n=43). This approach addresses limitations in existing methods by mitigating biases that arise from training on limited and artificial low-resolution datasets and providing a genome-wide representation of chromatin interactions. As a result, the model improves adaptability to real-world applications and enhances chromatin structure inference. To promote accessibility and further advance Hi-C data analysis, we have released both *TRUHiC* and *TRUHiC-LCL* as open-source frameworks, which surpass existing methods and enable broader applications in 3D genome research.

## 2 Results

In this study, we systematically evaluated *TRUHiC* across a wide spectrum of downsampled and experimental Hi-C datasets to establish its robustness and generalizability (Supplementary Table 1). Using the deeply sequenced GM12878 cell line, we first benchmarked *TRUHiC* on synthetically downsampled data (Subsections 2.1 and 2.2), demonstrating substantial improvements in both contact map reconstruction and structural feature recovery. We next applied *TRUHiC* to a low-resolution sample from the HGSVC2 cohort that was removed from our previous study^28^, GM12329, where *TRUHiC* enhancement resulted in a nearly 27-fold increase in the number of detectable chromatin loops compared to the original sample (Subsection 2.3). To further validate the performance of *TRUHiC* in a real experimental context, we compared enhancement results for NA19317 from the HGSVC3 cohort (∼8 kb resolution) against an independently generated ultra-high-resolution dataset from the same individual (GM19317, ∼3 kb resolution), observing marked improvements in concordance with the experimental ground truth (Subsection 2.4). We then systematically examined the performance of *TRUHiC* across varying levels of sequencing depth (Subsection 2.5), and extended the evaluation across additional cell types and species to confirm cross-context generalization (Subsection 2.6). In the end, we introduced a cell line-specific (CLS) variant of the model, *TRUHiC-LCL*, trained on 43 lymphoblastoid cell line datasets, which can be readily applied to analyze Hi-C data in various practical scenarios (Subsection 2.7).

### 2.1 Super-resolution reconstruction of Hi-C contact maps from downsampled low-resolution Hi-C data

*TRUHiC* takes low-resolution Hi-C contact maps as input and attempts to augment their resolution. The architecture of *TRUHiC* (see Figure 5) is based on the U-2 Net, and we equipped it with a transformer block to enable the resulting models to capture both short- and long-range interactions among the genomic regions and make context-aware predictions. We use low-resolution contact maps as the input and train the model in a supervised manner to predict the respective high-resolution contact maps. Also, we leverage mean absolute error (MAE) and signal-to-noise ratio (SNR) in our loss function with equal weights. To rigorously assess *TRUHiC*’s performance in reconstructing high-resolution Hi-C contact maps, we trained the model on chromosomes 1-17 of the GM12878 cell line and tested the resulting model on chromosomes 18-22 with a down-sampling rate of 1/16 (see Methods for downsampling details). For a comprehensive comparison with existing models, including *DFHiC*, *HiCNN2*, and *HiCARN*, we applied identical data preprocessing to that of *TRUHiC* to ensure the exact same data input is used to train and evaluate every method. Next, we saved each model for the subsequent experiments in this subsection and subsections 2.2 and 2.4. To comprehensively evaluate the quality of the enhanced Hi-C contact maps, we employed multiple image quality assessment metrics that capture different aspects of reconstruction fidelity (see Methods for details), including Peak Signal-to-Noise Ratio (PSNR), Signal-to-Noise Ratio (SNR), Spearman correlation coefficient (SPC), Pearson correlation coefficient (PCC), Structural Similarity Index Measure (SSIM), Mean Squared Error (MSE), Jaccard Index (JI), F1 scores, and GenomeDISCO Scores (GDS)^29^. These metrics were calculated to compare the enhanced low-resolution Hi-C matrices generated by *TRUHiC* against the competing methods. As summarized in Table 1, *TRUHiC* consistently outperforms the other enhancement methods across all metrics in the test set.

**Table 1.**
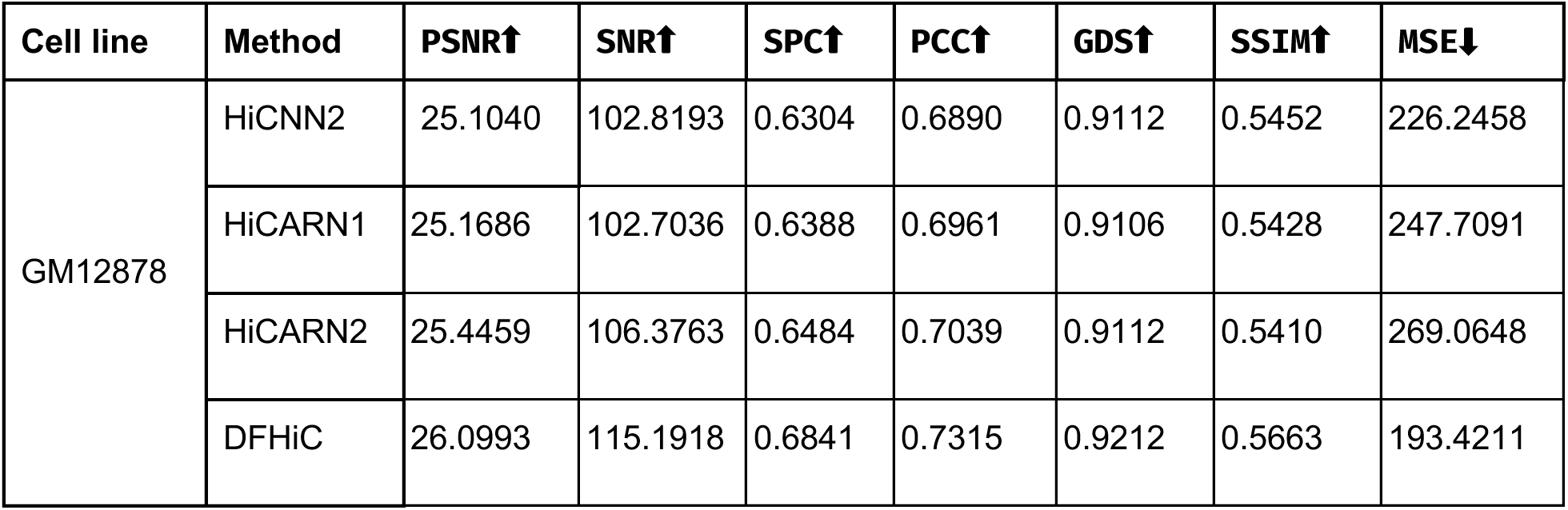
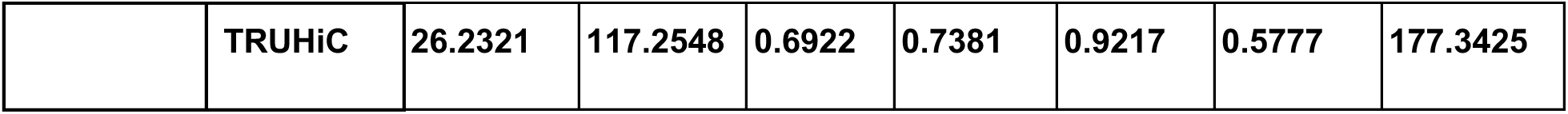
Comparison of vision metrics results averaged across test chromosomes (18-22) for each enhancement method. The arrows in each column indicate whether a higher (⬆) or lower (**⬇)** value is better, and the best score for each metric is bolded.

All enhancement models, including *TRUHiC*, were trained on an 80GB A100 Nvidia GPU, and the training time and memory consumption for each model on the Hi-C dataset with an input size of 40*40 are summarized in Supplementary Table S2. *TRUHiC* demonstrates a balanced trade-off between computational speed and memory efficiency compared to competing methods. *TRUHiC* does not require any additional data normalization or denormalization and uses the raw data directly. In contrast, *DeepHiC* and *HiCARN* need additional data normalization and denormalization procedures, as described in their papers. However, during experiments, we observed that *DeepHiC* and *HiCARN* performed better on raw data. Therefore, all competing models in this study were trained and evaluated on the raw Hi-C data to ensure a fair comparison.

### 2.2 Hi-C structural features reconstruction from downsampled low-resolution Hi-C data

To assess the effectiveness of *TRUHiC* in recognizing important 3D chromatin structures, namely TADs and loops, we employed two widely used tools: *Insulation scores (IS)*^30^ and *HiCCUPs*^31^. *IS* is primarily developed to detect TAD boundaries, with the regions between the two adjacent significant boundaries defined as TAD regions. *HiCCUPs* identifies chromatin loops by detecting enriched pixels, where contact frequencies within a pixel are compared to surrounding regions to determine significant looping interactions. We applied these methods to both high-resolution (HR) Hi-C datasets and enhanced contact maps generated by different models. To quantify the consistency of TAD predictions, we calculated the Jaccard Index (JI) and F1 score between the detected TAD boundaries in HR and enhanced datasets. The results for chromosomes 18-22 in the GM12878 cell line are summarized in Figure 1b-d and Table 2. The analysis reveals that *TRUHiC* consistently generates TAD boundaries with higher similarity to the high-resolution dataset compared to the competing methods. Unlike *HiCNN2* and *HiCARN*, *TRUHiC* does not overestimate the number of detected TAD boundaries, maintaining consistency with the HR dataset and indicating its effectiveness in preserving the integrity of chromatin structure.

**Table 2.** Comparison of TAD boundaries and chromatin loops detected by different models on test chromosomes (18-22) in the GM12878 cell line. The number shows the average value among the five test chromosomes. The arrows in each column indicate whether a higher (**⬆)** or lower value (**⬇)** is better, and the best-performing score in each category is highlighted in bold. Additionally, the higher validated loop anchor ratio in LR is attributed to the small number of loops detected in the LR data, where the majority of the identified loops successfully passed validation. The results for each test chromosome are summarized in Supplementary Tables S3-S6.

| Cell line | Method | TAD boundary<br>JI↑ | TAD boundary<br>F1↑ | Loop<br>JI↑ | Loop<br>F1↑ | Validated loop<br>anchor ratio<br>(%)↑ |
| --- | --- | --- | --- | --- | --- | --- |
|  | LR | 0.2771 | 0.4262 | 0.1564 | 0.2740 | 70.8700 |
|  | HiCNN2 | 0.2796 | 0.4304 | 0.2538 | 0.4415 | 68.5920 |
| GM12878 | HiCARN1 | 0.2870 | 0.4388 | 0.2654 | 0.4515 | 68.2800 |
|  | HiCARN2 | 0.3015 | 0.4568 | 0.2617 | 0.4446 | 72.8980 |
|  | DFHiC | 0.3324 | 0.4944 | 0.2775 | 0.4655 | 77.1280 |
|  | TRUHiC | **0.3479** | **0.5122** | **0.2992** | **0.4920** | **77.3360** |

We applied *HiCCUPs* to the original HR Hi-C matrices, downsampled low-resolution matrices, and enhanced-resolution matrices generated by *TRUHiC* and other competing methods to identify chromatin loops. We aimed to recover a higher number of true positive chromatin loops while maintaining structural integrity across test chromosomes. The results, presented in Figure 2, illustrate the number of loops identified by *HiCCUPs* for each method compared to the original loop calls in the HR matrices, with the overlapping sections indicating shared loop interactions. Additional results for other chromosomes can be found in Supplementary Figure S3. As expected, significantly fewer reliable loops can be detected from the low-resolution Hi-C data. Notably, *TRUHiC* outperformed competing methods across all test chromosomes by retaining the highest ratio of recovered significant chromatin loops over identified spurious loops.

**Figure 2.**
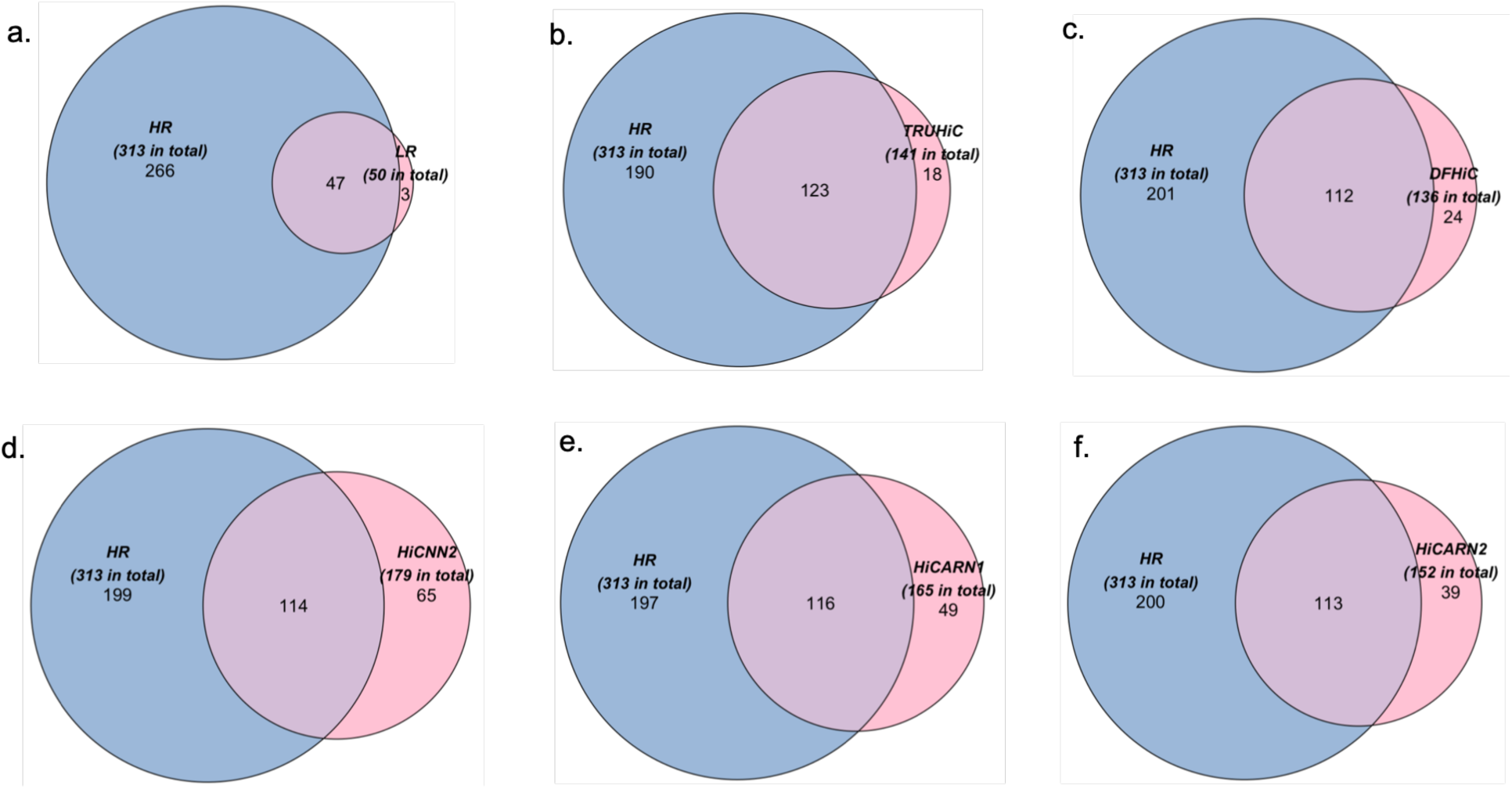
Comparison of chromatin loops detected on chromosome 20 of the GM12878 cell line. The numbers of chromatin loops are obtained by running *HiCCUPs* on the Hi-C data recovered by different enhancement methods from down-sampled low-resolution Hi-C data. Each blue circle represents the total number of loops detected in the high-resolution (HR) Hi-C datasets, while the corresponding pink circle represents the loops identified on the HiC data from the downsampled sample (Figure 2a) and on the enhanced Hi-C data using each method (Figures 2b-2f). The overlapping section indicates the intersections of blue and pink circles and thus represents loops that are true positives resulting from each enhancement method. The results for other cell lines are shown in Supplementary Figure S4.

We calculated the Jaccard Index (JI) and F1 score to assess the consistency of the TAD boundary and loop calls. Specifically, the JI measures the similarity between two sets by calculating the intersection ratio counting (at least one bp overlap) and the union, using the following formula:

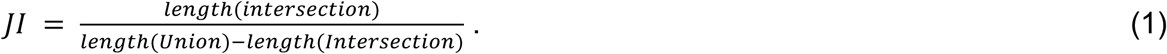

For TAD boundaries, we considered two regions identical if they shared at least a one base pair (bp) overlap. For loop calls, we defined two loops as matching if their positions fell within the range of +/− 5 kb. The F1 score was computed as follows^32^:

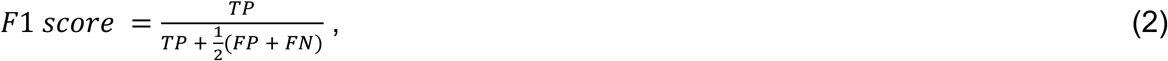

where we defined true positives (TP) as predicted loops that fall within the spatial range of +/− 5 kb around the loops identified in the ground truth. False positives (FP) refer to predicted loops that fall outside this flanking window. False negatives (FN) are ground-truth loops that lack a corresponding predicted match within the same range. True negatives were not considered in the calculation, as they represent the majority of the genomic space and provide limited meaningful information.

We further evaluate the quality of loop calling for each method by calculating the proportion of CCCTC-binding factor (CTCF) validated loop anchors (see Methods for details). Prior studies have shown that the majority of loop anchor loci are bound by the insulator protein CTCF, along with cohesin subunits RAD21 and SMC3^3^. Therefore, we expect that our model will be able to predict a higher proportion of loop anchors that can be validated as CTCF-supported loop anchors co-occurring with CTCF, RAD21, and SMC3 ChIP-seq peaks. We applied this to the LR, *TRUHiC,* and other competing methods and reported the validated loop ratio in Table 2. Note that the comparative JI and F1 scores of the TAD boundary for LR align with the theoretical basis of the IS algorithm, which was initially designed for detecting TAD features in low-resolution Hi-C data. While IS demonstrates relative robustness across different data resolutions, our method achieves discernible improvements across all aspects, further reinforcing its effectiveness.

During the experiments, we questioned the extent to which vision metrics correlate with biologically meaningful feature metrics. While prior studies have reported visual metrics, we did not find compelling discussions on why such metrics are specifically important for the Hi-C enhancement task. Commonly used vision metrics such as SPC, PCC, SNR, SSIM, and PSNR are widely employed for image quality assessment; however, their direct relevance to biological features, such as chromatin loops and TADs, remains unexplored. To investigate this, we computed 𝑅^7^ scores between various visual metrics and biological feature metrics we obtained from the above experiments to determine which visual assessments best reflect biologically significant structures in Hi-C data (Figure 3).

**Figure 3.**
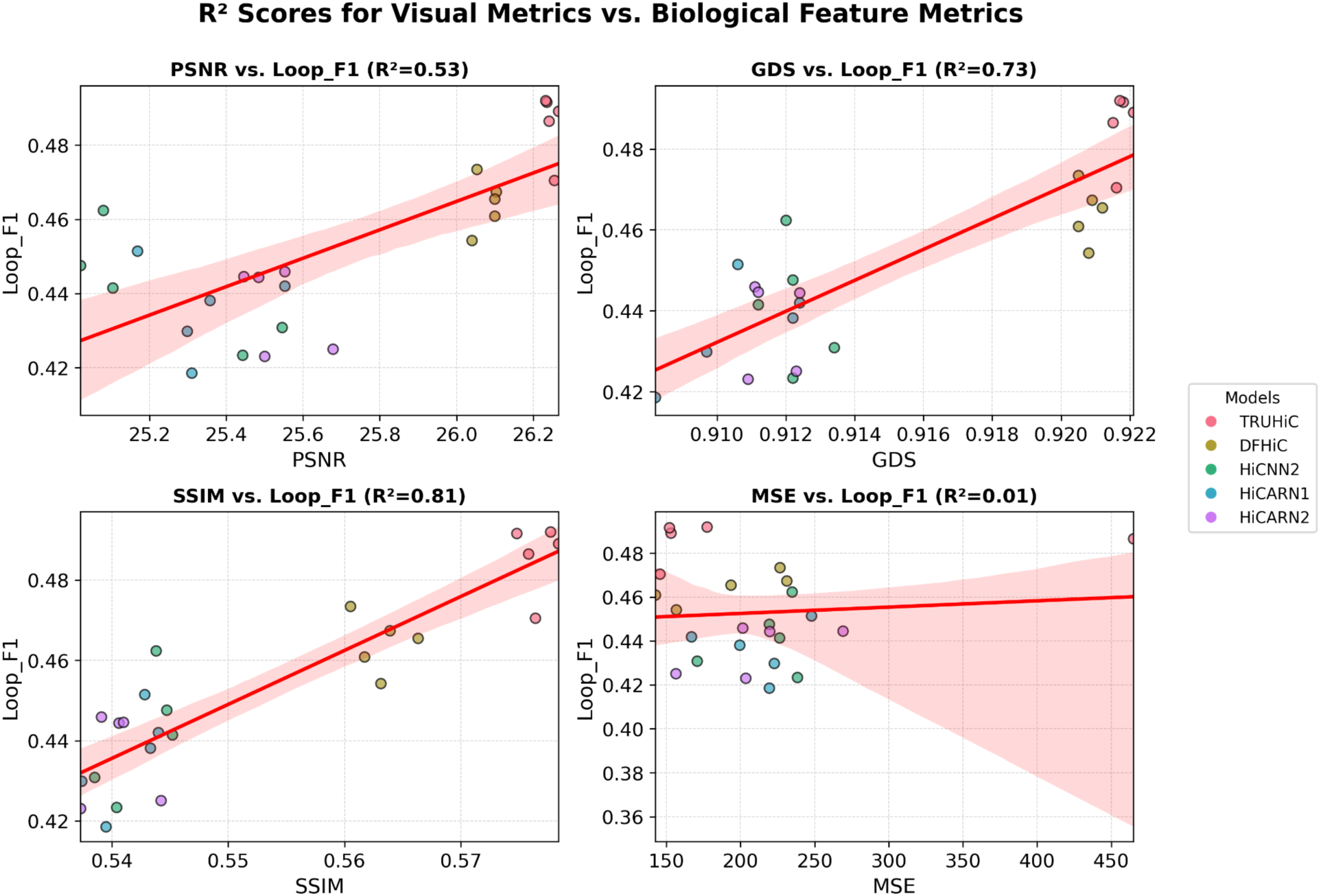
Correlations between vision quality metrics and biological feature metrics. The plots display 𝑅^7^ scores quantifying the relationships between selected vision metrics on the X-axis (PSNR, GDS, SSIM, and MSE) and biological feature matrices on the Y-axis (F1 score) for chromatin loops. The full results are shown in Supplementary Figure S5.

Our analysis revealed that PSNR, SNR, SPC, PCC, GDS, and SSIM exhibit strong correlations with TAD boundary feature metrics (𝑅^7^ scores 0.85-0.88), suggesting that these visual measures effectively benefit hierarchical chromatin organization identification. Likewise, SSIM and GDS correlated well with loop-based metrics (𝑅^7^ scores 0.73-0.81), indicating their utility in evaluating fine-scale structural features. In contrast, PSNR, SNR, SPC, and PCC showed only moderate correlations with loop-based metrics (𝑅^7^ scores 0.53-0.71), whereas MSE showed no association with any biological metrics (𝑅^7^ scores = 0.01), underscoring its limitations in assessing structural fidelity. These findings underscore the importance of selecting biologically relevant evaluation metrics when developing and benchmarking Hi-C data enhancement models.

### 2.3 Enhancing resolution in experimentally sparse Hi-C data

We applied our pre-trained *TRUHiC* model, trained on GM12878 with a 1/16 downsampled rate, to augment an actual low-resolution Hi-C data set on GM12329, obtained from HGSVC2, with an original contact map resolution of approximately 18 kb. This sample of GM12329 had been excluded from our previous research study^28^ due to its low sequencing quality, which resulted in the detection of only 158 chromatin loops at 10 kb resolution. To comprehensively evaluate its performance, we assessed all 3D genome structure features and all the reproducibility scores of GM12329 at a genome-wide scale. After enhancement with *TRUHiC*, we identified 4,177 chromatin loops across all autosomes, with all 158 loops originally detected in the data also present (Supplementary Figure S6). We further compared the *TRUHiC*-enhanced TAD boundaries in GM2329 with the recently released integrative TAD catalog in lymphoblastoid cell lines^28^. The results of this experiment, presented in Supplementary Table S7, showcase that *TRUHiC* outperforms the competing methods in accurately identifying loops and TAD boundaries in real-world Hi-C data as well. However, we observed a decline in performance across all evaluation metrics for GM12329 compared to our results on the downsampled GM12878 dataset (Sections 2.1 and 2.2). This finding is consistent with a recent study assessing the generalizability of deep learning-based Hi-C resolution improvement methods, which reported that existing deep learning approaches struggle to generalize to experimentally derived sparse Hi-C datasets, with performance reductions of up to 57%^27^. These results highlight a critical limitation in current deep learning frameworks and underscore the need for improved strategies to enhance model generalizability. While our proposed *TRUHiC* framework outperforms existing methods, addressing its robustness on real sparse data remains an important direction for further investigation, as discussed in which we investigate further in Subsection 2.6 (CLS model).

### 2.4 Enhancement validation through experimentally high-resolution data

To further validate *TRUHiC* in a real experimental setting, we analyzed the NA19317 Hi-C dataset from the HGSVC3 cohort, which was generated at a sequencing resolution of approximately 8 kb. We benchmarked the enhancement results resolved from *TRUHiC* against an independently generated ultra-high-resolution Hi-C dataset from the same individual (GM19317), with a resolution of roughly 3 kb. This setup provided a unique opportunity to directly assess model performance against a true high-resolution experimental reference. When applied to the HGSVC3 NA19317 data, *TRUHiC* increased the total number of detectable chromatin loops by more than tenfold at 10 kb resolution (from 1,210 to 15,926), underscoring its ability to recover focal interaction features that were largely missed in the raw data. Furthermore, benchmarking against the ultra-high-resolution GM19317 reference revealed that both *TRUHiC* and the competing models improved upon the original HGSVC3 NA19317 feature calls in terms of Jaccard Index and F1 scores. Among these, *TRUHiC* achieved the strongest performance, yielding the highest Jaccard Index and F1 scores across both TAD boundary and chromatin loop identification, as well as superior results on image-based evaluation metrics (Supplementary Table S8). These findings demonstrate that *TRUHiC* not only preserves the integrity of feature detection in datasets generated at moderate sequencing depth, but also refines boundary placement and loop anchoring in a manner that more faithfully reflects ultra-high-resolution experimental data. Collectively, this establishes *TRUHiC* as a robust framework for extracting more precise 3D genome features and extending the scientific value of existing datasets, where deeper coverage may not be feasible.

### 2.5 Performance assessment at different levels of data sparsity

To evaluate the robustness of our method under varying levels of data sparsity, we extended our analysis beyond the 1/16 downsampled dataset by randomly down-sampling the high-resolution (10 kb) GM12878 cell line Hi-C data. We used 1/50 and 1/100 downsampling ratios, resulting in progressively lower resolution datasets (500 kb and 1 Mb, respectively), allowing us to assess *TRUHiC*’s performance relative to competing methods under different sparsity conditions. All models were retrained separately for each downsampled dataset and subsequently tested on chromosomes 18-22 using the same evaluation metrics: PSNR, SNR, SPC, PCC, GDS, SSIM, MSE, and biologically relevant metrics for TAD boundary and loop detection. The results, summarized in Supplementary Tables S9 and S10, demonstrate that *TRUHiC* predominantly outperforms other models at down-sampling rates of 1/50 and 1/100. As expected, increasing the down-sampling rate led to a decline in enhancement performance across all models due to the greater loss of structural information at higher sparsity levels.

### 2.6 Generalization across different cell types and species

We aimed to assess *TRUHiC*’s capability in enhancing low-resolution Hi-C data across multiple human cell lines (K562, IMR90, and NHEK), as well as a mouse cell line (CH12-LX), to benchmark model generalization across different cell types and species. These datasets were originally processed at the same resolution as GM12878 (10 kb) and were subsequently downsampled by 1/16 to a 160 kb resolution. In this experiment, the pre-trained GM12878 models from Subsection 2.1 were directly applied to enhance different down-sampled Hi-C data from the three additional human cell lines and the mouse line. The test chromosomes for the human cell lines remained the same as GM12878 (18-22), while chromosomes 16-19 were selected as the test set for the mouse cell line. To evaluate the models’ effectiveness, we used the previously established evaluation metrics, which are outlined in Supplementary Table S11. We observed that *TRUHiC* achieved higher performance scores compared to the competing methods in three of these cell lines (K562, IMR90, and CH12-LX) while maintaining competitiveness in the remaining human cell line (NHEK).

To further investigate the capability of *TRUHiC* to identify TADs and loops, we employed the same TAD and loop callers, *IS* and *HiCCUPs,* to detect these two features on the test chromosomes across different cell lines. As shown in Supplementary Table S12, *TRUHiC* achieved higher JI values, indicating a greater number of true positive TAD boundaries and loops while maintaining consistently lower false positive rates across all cell lines and species compared to other methods. These results demonstrate *TRUHiC*’s comparative generalization power across different cell lines and species.

### 2.7 Towards robust Hi-C data enhancement using a cell line-specific (CLS) model

Beyond our primary experiment using a 1/16 downsampled ratio, we generated four additional replicates of data with the same 1/16 downsampling ratio using different random seeds, constituting a total of five low-resolution datasets (Supplementary Figure S7) to try to mimic more diverse data distribution observed in real-world Hi-C experiments. We trained *TRUHiC* and other competing models on each of these five datasets separately (the primary dataset plus four replicates) and observed inconsistent prediction performance across different replicates and methods (Supplementary Tables S13 and S14). We argue that these fluctuations in model performance arise from inherent variations in the input Hi-C data distributions. Statistical analysis supports this hypothesis, as indicated by a significant Kruskal-Wallis Test *p-value* (< 0.05) followed by a Kolmogorov-Smirnov Test p-value for pairwise comparisons (Supplementary Table S15). To address this issue, we propose a distinctive model training strategy that integrates a diverse set of real Hi-C data rather than relying solely on a single downsampled sample. This approach is inspired by recent advancements in foundational models, which have demonstrated the effectiveness of large-scale data aggregation in various domains, such as natural language processing^33–37^ and computational biology^38–44^.

Lymphoblastoid cell lines (LCLs) are widely studied in large-scale genomic research and are of particular importance for functional genomic and pharmacogenetics studies in humans^45–48^. As a model system, LCLs enable scientists to study gene regulation, genetic variation, and disease mechanisms at the population level using a consistent cell type that can be easily expanded from small blood samples. Given their significance, a cell line-specific (CLS) model tailored for enhancing Hi-C data in LCLs is of great importance to the research community. Despite the merits a foundational model trained on diverse species and cell lines presents, a CLS model is better suited to capture the unified chromatin interaction patterns specific to the same cell line^49,50^. That is, a CLS model is inherently biased toward the distinct structural patterns unique to a specific cell line, which could be otherwise lost in favor of more common patterns across species and cell lines in a foundational model.

Instead of pooling Hi-C data at the matrix level, as done in Li et al.’s study^28^, our proposed CLS modeling paradigm learns hierarchical chromatin interaction patterns from multiple independent biological samples during training, enabling it to generalize across diverse Hi-C datasets. Our CLS model framework is built upon the *TRUHiC* architecture and is trained on a large-scale dataset of HGSVC Hi-C data from Li et al.’s study^28^ (Figure 4a). To systematically assess model scalability, we designed two hierarchical training sets: a small dataset consisting of 10 unique HGSVC biological samples (training set S) and another large dataset including an expanded set of 43 HGSVC biological samples, with GM12878 excluded (training set L). Both models were evaluated using a test set comprising three biological sample data (GM11168, GM13977, and GM18951) from Harris et al.’s study^51^, ensuring that none of these samples were included in any of the training sets.

**Figure 4.**
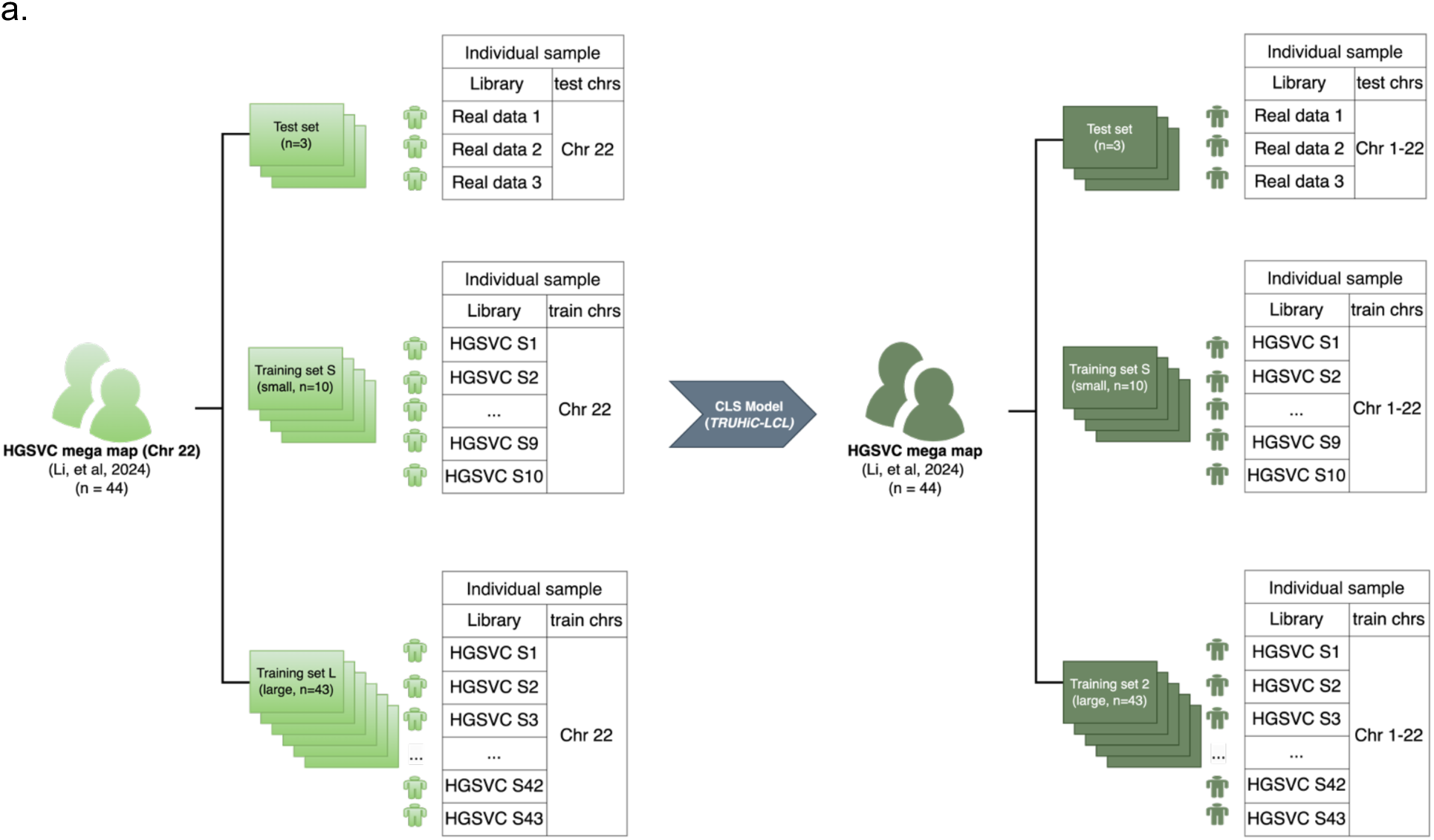
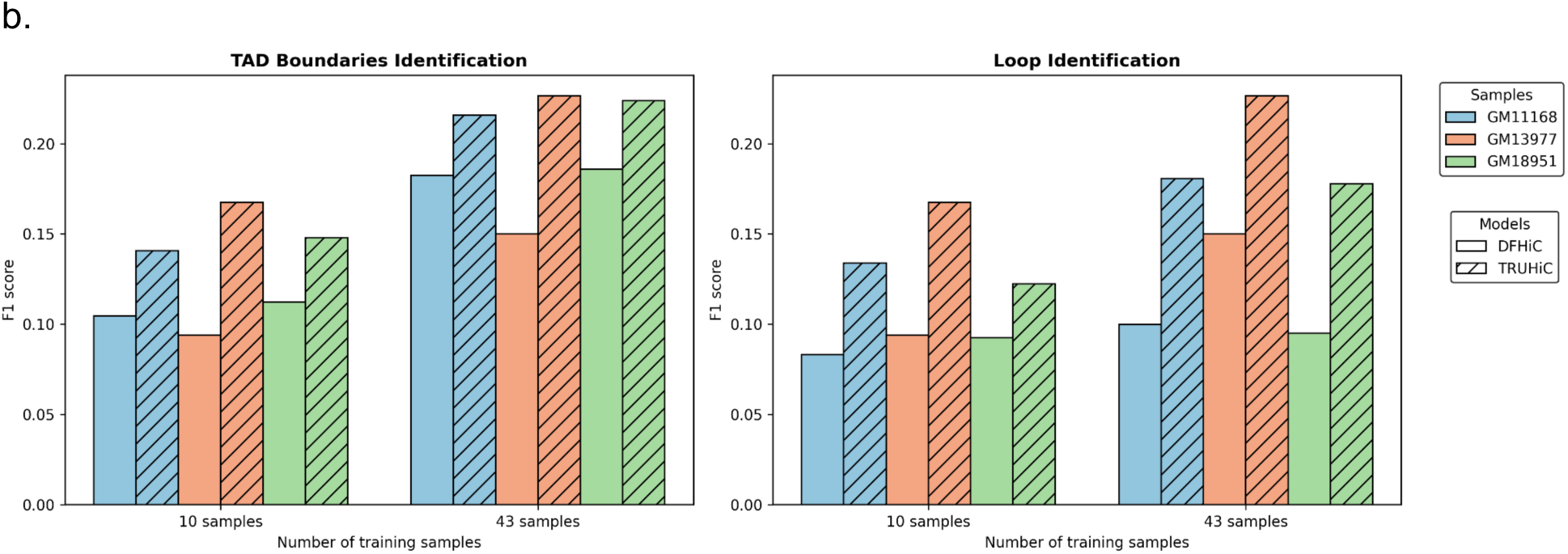
The training strategy of building an LCL cell line-specific (CLS) model and performance comparison of *TRUHiC-LCL* and *DFHiC-LCL* on chromosome 22. **a).** This schematic illustrates our proposed approach for developing a Hi-C CLS model (*TRUHiC-LCL*) by aggregating diverse real Hi-C datasets from HGSVC (Li et al., 2024, n=43). We train an LCL-model with two hierarchical training sets: Training Set S (small, n=10) and Training Set L (large, n=43, GM12878 is excluded). The left panel represents training using chromosome 22, while the right panel shows the potential to extend this framework to whole-genome training (chromosomes 1-22), demonstrating the scalability of *TRUHiC-LCL* for constructing a Hi-C foundation model that enhances data consistency and generalizability across multiple biological samples. **b).** The figure presents F1 scores for TAD boundary and loop identification using a small training set S (10 samples) and a large training set L (43 samples) across three real Hi-C samples (GM11168, GM13977, and GM18951). Performance is compared between the state-of-the-art *DFHiC* model and our *TRUHiC* framework. *TRUHiC* consistently outperformed *DFHiC* across both training sample sizes and demonstrated further improvements as the training set increased.

Due to computational constraints, the initial model training was conducted on chromosome 22. As shown in Supplementary Tables S16, S17 and Figure 4b, the model trained on training set L yielded significantly improved performance across all vision and biological feature evaluation metrics compared to the model trained on smaller training set S. These findings support our hypothesis regarding the effectiveness of our CLS-model framework, *TRUHiC-LCL*, demonstrating that a CLS-model trained on the whole genome Hi-C data could even more effectively capture complex chromatin interaction patterns present in real Hi-C datasets, surpassing the limitations of single-sample training approaches. In this study, we release the pre-trained chromosome 22 *TRUHiC-LCL* model trained on the training set L with 43 Hi-C samples, providing an open-access resource for future applications in Hi-C data enhancement and chromatin structure characterization.

## 3 Discussion

Recent advances in chromosome conformation capture technologies like Hi-C have provided critical insights into chromatin folding and genome organization. However, the resolution of Hi-C data is often constrained by sequencing depth and experimental limitations, making it challenging to accurately detect TADs and chromatin loops. To address this, we proposed *TRUHiC*, a computational framework designed to significantly enhance low-resolution Hi-C data with high fidelity. The gains in resolution allow for more precise identification of higher-order chromatin structures, enabling a deeper investigation into genome organization. Such improvements can facilitate studies on the spatial relationships between regulatory elements and their target genes, help identify structural variations that impact gene expression, and ultimately provide new insights into how chromatin architecture shapes gene regulation and cellular function.

Expanding on this, we introduced *TRUHiC-LCL*, a lymphoblastoid cell line (LCL) specific model trained on enriched Hi-C datasets aimed at improving generalizability and robustness across different sequencing depths and experimental conditions for LCL Hi-C data. We provide a pre-trained *TRUHiC-LCL* model based on HGSVC data, specifically for chromosome 22 of LCLs, developed within our constraints of computational limitations. We anticipate that these models will be further enhanced with regard to their robustness and performance in the future by incorporating additional LCL Hi-C data from consortia such as HGSVC^52^, HPRC^53^, and the 4D nucleome project^54^, as well as studies like Harris et al.^51^. Moreover, our open-access framework is adaptable and enables researchers to train and finetune their own CLS models and extend the approach to whole-genome Hi-C data in various cell lines, tissues, and species, facilitating broader applications in chromatin structure analysis.

Our experimental results support the idea and demonstrate the feasibility of building a foundational model for Hi-C data using our approach. By leveraging existing Hi-C datasets, this method enables the enhancement of low-resolution data, providing a scalable framework for understanding chromatin architecture in species with limited genomic resources. We anticipate that the approaches demonstrated in this study will be transferable to future extensions to build foundation models with massive Hi-C data when available, with abundant opportunities to be applied to various applications across diverse organisms other than human cell lines, such as plants and agriculturally important species like soybeans. However, significant challenges remain in the development of such generalized models that await future explorations.

One key limitation is the availability of high-quality, diverse Hi-C datasets for training. Currently, *TRUHiC* has been primarily trained on GM12878, a well-characterized human cell line. Although *TRUHiC-LCL* has been trained on a large scale of LCL Hi-C data, the reliance introduces a potential bias in performance, as the model may become overly tailored to the specific chromatin features of the dataset of that single cell line, limiting its generalizability. To improve model robustness, the inclusion of Hi-C data from additional cell types and species is critical. A more diverse training set would reduce bias and enhance the model’s ability to generalize across different chromatin architectures, such as those found in plant genomes or organisms with unique genomic configurations.

The performance discrepancy observed when applying the model to different datasets further underscores this limitation. A biased training dataset may result in reduced accuracy when used on species or conditions with chromatin features that deviate significantly from those of GM12878. Overcoming this challenge requires efforts to collect more Hi-C data, particularly high-resolution datasets from diverse backgrounds, to create a more comprehensive training set across different biological contexts. Future work should thus focus on expanding training datasets in both diversity and volume to reduce bias and enhance the model’s generalizability. Additionally, it is essential to explore how the model performs across datasets with varying resolutions and experimental protocols to help identify and address potential discrepancies. Coupling Hi-C data with complementary multi-omics datasets, such as ATAC-seq, ChIP-seq, and RNA-seq, could also enhance the model’s ability to link chromatin structure with gene regulation and functional outcomes, providing deeper insights into genome organization and transcriptional regulation. With the continued expansion of training datasets and continued model refinement, this method has the potential to advance our understanding of chromatin architecture across a wide range of organisms and biological contexts. Ultimately, it could contribute to accurate reconstructions of three-dimensional genome organization, facilitating new insights into gene regulation, epigenetics, and genome function.

## 4 Materials and methods

### 4.1 Materials

We utilized a published high-resolution Hi-C dataset of human cell type GM12878, three additional different human cell types, K562, NHEK, and IMR90, and one mouse cell type CH12-LX from the GEO database (accession number GSE63525)^3^ and a recent study that integrated human cell types from 44 individuals^28^. To generate low-resolution Hi-C data, we applied a random down-sampling approach to the raw sequencing reads of each cell line using down-sampling rates ranging from 1/16 to 1/100, with the primary sampling rate set as 1/16. We trained our model using both high-resolution and low-resolution Hi-C data. These diverse sampling rates enable us to assess the model’s performance across different levels of raw reads sequencing depth. Both high-resolution and the corresponding low-resolution Hi-C matrices were partitioned into 40 × 40 non-overlapping blocks of 10 kb resolution with no normalization method applied (normalization set to NONE). We followed established practices in Hi-C super-resolution methodologies to preserve only small fragments where the genomic distance between two loci is < 2Mb, considering the typical average genomic distance of TADs to be < 1 Mb^5,6,13,14,16^. Chromosomes 1-17 comprised our training set, and chromosomes 18-22 constituted our test set. We split the training set into training and validation data following a 9.5:0.5 ratio during the training process of our model and all competing methods to have an identical and fair training regiment.

### 4.2 *TRUHiC* architecture

*TRUHiC* is a computational framework designed to enhance the resolution of Hi-C matrices through supervised training. The overall architecture of *TRUHiC* is portrayed in Figure 5. The backbone of the *TRUHiC* is a U-2 Net architecture, the successor of the U-Net architecture. A U-Net^55^ is an auto-encoder based model that has skip connections from the encoder layers to the respective decoder layers. A U-2 Net^25^ is a U-Net in which layers are replaced by U-Net-like blocks, termed ReSidual U-blocks (RSU). More specifically, a U-2 Net uses RSU-L and RSU-4F blocks. RSU-L blocks use a CNN-based symmetric U-Net architecture with L-1 encoder/decoder convolutional layers in addition to the pooling and upsampling operations. RSU-4F blocks are used as the bridge of the U-2 Net, as well as pre- and post-bridge blocks, using dilated convolutions. In the originally proposed U-2 Net framework, the encoder/decoder has four RSU-L blocks and one RSU-F block, and another RSU-F block is used as the bridging block between the encoder and the decoder. Another characteristic of the U-2 Net architecture is the presence of auxiliary outputs from the bridging block and each decoder block.

We customized the U-2 Net architecture for the Hi-C enhancement task by refining several key components to optimize performance and computational efficiency. At a high level, we reduced the number of encoder/decoder RSU-L blocks to two, which simplifies the network while retaining sufficient capacity to extract essential features. In our design, the encoder/decoder blocks play a critical role in capturing multi-scale information and reducing their number, which lowers computational complexity without significantly compromising performance. Additionally, we introduced a direct skip connection from the input to the outputs, inspired by the *DFHiC* model^13^. This connection helps preserve pixel-level details. To further improve feature representation, we replaced the traditional RSU-4F bridging block with a customized multi-head self-attention gating mechanism^56^ embedded within a transformer block (Supplementary Figure S8). This self-attention module mainly regulates channel-wise information through fully connected layers, thereby enhancing the quality of the generated feature maps. We also applied weighting to each auxiliary output. Specifically, the loss from each auxiliary output is multiplied by *1/e^x^*, where *x* is the index of the auxiliary output, starting at one for the final RSU-L block and ending at four for the transformer block. For example, the auxiliary output from the RSU-4F block is weighted by 1/e^3^. This approach prevents earlier layers from being forced to generate a fully refined prediction and instead allows them to reinforce the final output effectively. Finally, to preserve spatial resolution, a critical aspect for accurate Hi-C data enhancement, we removed the maximum pooling and upsampling operations from the RSU-L blocks (Supplementary Figure S8). Furthermore, we applied both L1 and L2 regularizations to the convolution layers to mitigate overfitting and improve the model’s generalization capabilities.

**Figure 5.**
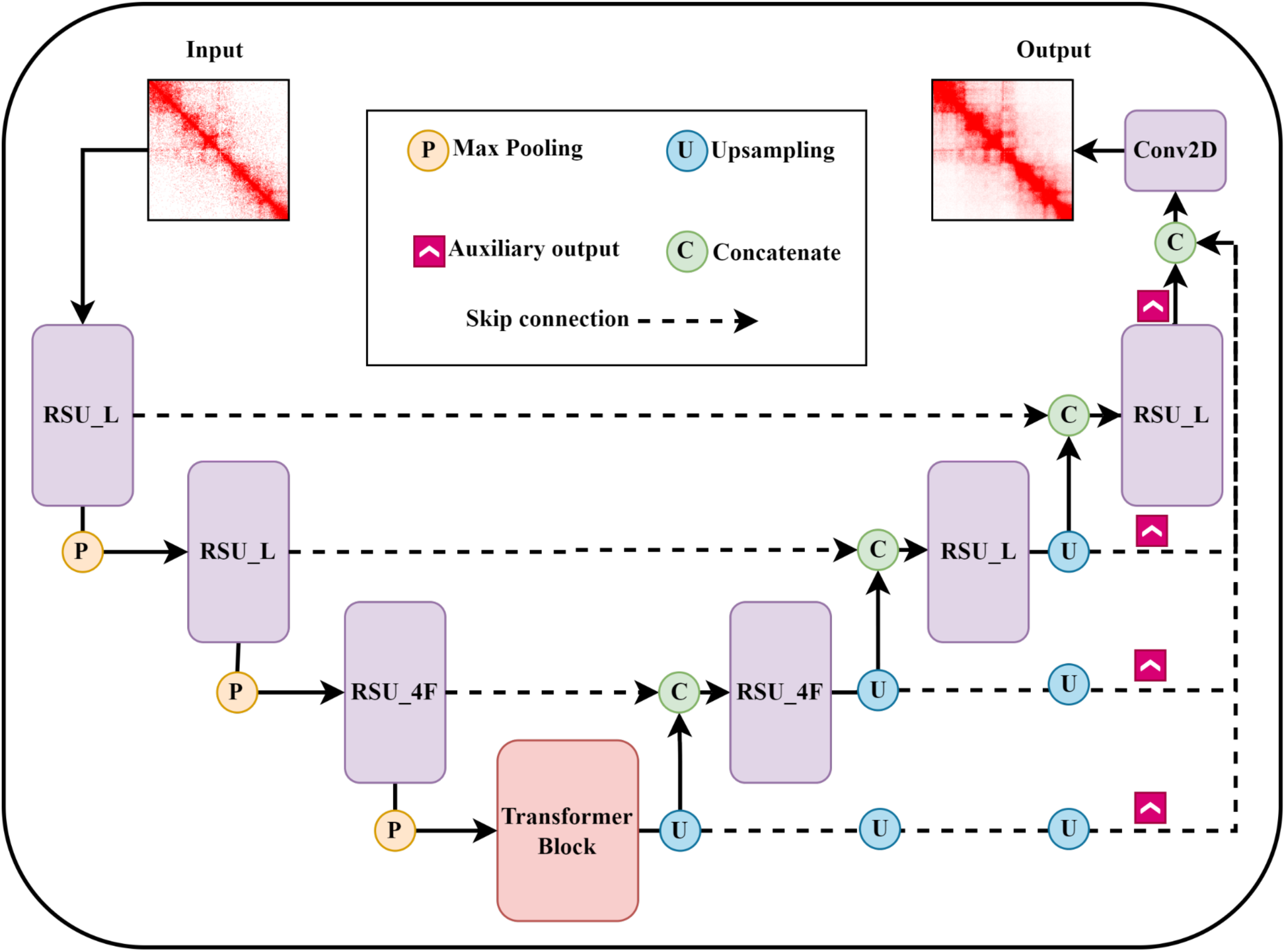
Overview of the *TRUHiC* architecture. We customized the U-2 Net architecture in several ways. A skip connection was added from the input to the output of the model. We replaced the U-Net bridge block in the model with (customized) multi-head self-attention gating wrapped up in a transformer block. We also removed MaxPooling and UpSampling layers to prevent the loss of information due to data compression. Additionally, auxiliary outputs have a weighted loss contribution. *TRUHiC* takes the low resolution of the Hi-C contact map as input and generates the super-resolution Hi-C contact map as output. A detailed illustration of RSU_L, RSU_4F, and Transformer blocks can be found in Supplementary Figure S8.

Additionally, we altered the loss function to include two terms as follows:

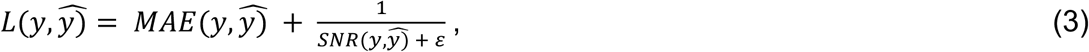

where MAE denotes the mean absolute error, and SNR is the signal-to-noise ratio, defined in Equation (6). We use the LAMB optimizer^57^ and early stopping. Also, we implement a learning rate reduction strategy upon a loss plateau to enhance convergence stability.

### 4.3 Identification of TAD boundaries and loops

We used the *Insulation Score* (*IS*) method to identify TAD boundaries in this study, which was originally designed to detect TAD boundaries and quantify the boundary strength of Hi-C data with limited resolution. For all experiments with GM12878 data, the Hi-C data were mapped to the hg19 human reference genome, and the KR normalized contact matrix at 10 kb resolution was used to compute insulation scores and boundary scores (BS). TAD boundaries were identified using the *FAN-C* toolkit (version 0.9.26b2) with a minimum boundary score cut-off value of 0.20, specifying a 100 kb window size, as referenced in the 4DN domain calling protocol^54,58^. For the experiments of HGSVC2 sample GM12329, and HGSVC3 sample NA19317, we used the hg38 human reference genome and applied SCALE normalization to the predicted contact map at 10 kb resolution, whereas the ultra-high-resolution sample GM19317 sample was detected at 5 kb resolution. For the *TRUHiC-LCL* model, we selected the 5 kb resolution to be consistent with the protocol of the integrative TAD catalog described in Li et al.’s study^28^.

The *IS* method operates by defining a sliding window along the diagonal of the Hi-C matrix and summing contacts within this window. Regions with low insulation scores (corresponding to high boundary scores) act as insulating boundaries and are identified as TAD boundaries. In contrast, regions with high insulation scores (low boundary scores) typically fall within TAD domains and are referred to as TAD regions, which represent the genomic intervals between adjacent TAD boundaries in this study. TADs with a size larger than 2 Mb were excluded from the analysis, and sex chromosomes X and Y were removed from all analyses due to sex-based variability in the samples. *Juicebox* software and the *FAN-C* toolkit in Python 3.7 were used to visualize insulation scores, TAD boundaries, and respective boundary scores.

Chromatin loops (and loop anchors) in the experimental samples GM12878, GM12329, and NA19317 were identified using *HiCCUPS* (GPU) at 10 kb resolution, whereas the ultra-high-resolution GM19317 sample was detected at combined 5 kb and 10 kb resolution. For the three real samples enhanced by the *TRUHiC-LCL* model, loop calling was performed at 5 kb resolution. The data alignment and matrix normalization procedures for the experiments with the GM12878 sample, actual samples, and *TRUHiC-LCL* followed the same approach as described in TAD boundary identification. The Jaccard Index was computed using the *bedtools jaccard* command, and the F1 score was calculated based on the equation provided in the previous section and implemented using our custom Python script.

### 4.4 CTCF loop anchor validation with ChIP-seq datasets

ChIP-Seq experimental datasets for CTCF, RAD21, and SMC3 for each cell line were obtained from Rao et al.’s study^3^. For each loop anchor, we expanded its region by ±5 kb flanking windows and merged overlapping or adjacent intervals into a single larger interval. A loop anchor was classified as CTCF-supported if its expanded regions fully contained CTCF ChIP-Seq peak, RAD21 ChIP-Seq peak, and SMC3 ChIP-Seq peak simultaneously. In cases where RAD21 or SMC3 ChIP-Seq data were unavailable for a given cell type, a loop anchor was considered CTCF-supported if the expanded anchor overlapped with CTCF and either SMC3 or RAD21 peaks. If only CTCF ChIP-Seq data were available, the loop anchor was required to show a direct overlap with a CTCF ChIP-Seq peak to be classified as CTCF-supported. The validated loop anchor ratio was calculated as the percentage of CTCF-supported loop anchors among the total unique loop anchors. We reported this value as a matrix to evaluate the accuracy of loop calling across different methods. The validation experiment was not performed for the CH12-LX cell line due to the absence of corresponding ChIP-Seq datasets provided in Rao et al.’s study. A detailed list of CTCF ChIP-Seq datasets used in this analysis is provided in Supplementary Table S18.

### 4.5 Baseline Models

We selected *HiCNN2*, *HiCARN1*, *HiCARN2*, and *DFHiC* methods for benchmarking our proposed method. While pre-trained model weights for a number of the mentioned methods are publicly available, we opted to train them all from scratch to ensure fair and consistent evaluations. Each model was implemented using its official source code to maintain fidelity to the original methods. In the case of *DFHiC*, we re-implemented the code in Tensorflow 2.14 and added improvements to it for scheduling the learning rate to prevent premature loss convergence. For the *HiCNN2* and *HiCARN* (*HiCARN1* and *HiCARN2*) methods, we used the source PyTorch implementations without any major changes other than cleaning up the code and adding command-line arguments for ease of training and inference. We are providing the implementations of these models on our repository at https://github.com/shilab/TRUHiC for improved reproducibility of the results.

### 4.6 Evaluation metrics

To assess the performance of our model and the quality of the generated enhanced Hi-C samples, we considered the output as an image and employed various evaluation metrics, which include Mean Squared Error (MSE), Structural Similarity Index (SSIM), Peak Signal-to-Noise Ratio (PSNR), Signal-to-Noise Ratio (SNR), Spearman Correlation Coefficient (SPC), Pearson Correlation Coefficient (PCC) and GenomeDISCO Scores (GDS).

Firstly, MSE is used to calculate the average squared difference between the model-predicted Hi-C matrix and the real high-resolution Hi-C matrix, which can effectively capture the average discrepancy at the pixel level. SSIM score evaluates the structural similarity between the enhanced output and the ground truth Hi-C matrix, with higher scores indicating greater preservation of structural integrity. Furthermore, we used both PSNR and SNR to measure the quality of the reconstructed Hi-C contact maps. Specifically, SNR quantifies the ratio of signals relative to background noise, whereas PSNR measures the ratio of the maximum possible signal power to the power of corrupting noise in the enhanced contact matrix and the target real high-resolution matrix. The higher both values are, the more unwanted noise is removed. PSNR and SNR are formulated in Equations (5 and 6), respectively. We employ SPC and PCC to evaluate the correlation between the predicted Hi-C matrix and the actual Hi-C matrix along the matrix diagonal. Additionally, we used a concordance measure named GenomeDISCO, which was developed to assess the similarity between a pair of contact maps received from 3C experiments. We provide the equations for these metrics as follows^16^:

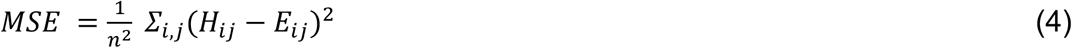

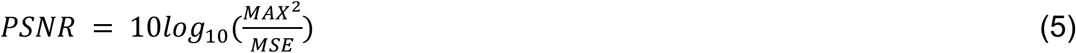

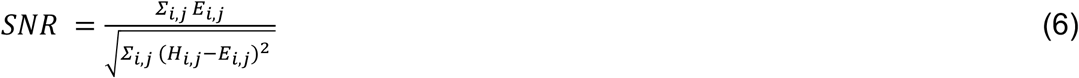

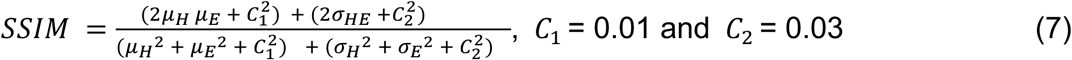

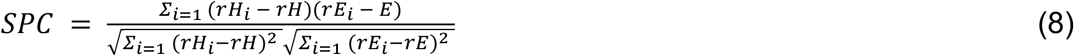

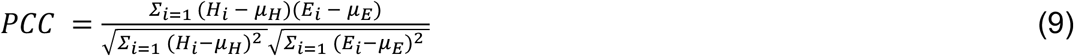

where 𝐻*_ij_* is the pixel in the real HR Hi-C matrix, and 𝐸*_ij_* is the pixel in the enhanced Hi-C matrix. MAX denotes the maximum possible value in samples, while μ and 𝜎 correspond to the mean and variance, respectively. 𝜎*_HE_* is the covariance of 𝐻 and 𝐸, and 𝑟 is the rank. Mean, variance, and covariance in the SSIM formula are calculated using a Gaussian filter, and C_1_ and C_2_ are constants used to stabilize the calculations. We used the SSIM implementation provided in *DeepHiC*.

### 4.7 Data and Code Availability

The GM12878, K562, IMR90, NHEK, and CH12-LX datasets supporting this study are publicly accessible in the GEO database under accession number GSE63525, available at https://www.ncbi.nlm.nih.gov/geo/query/acc.cgi?acc=GSE63525. The raw sequencing Hi-C data generated by HGSVC2 discussed in this study can be downloaded directly at the following link: https://ftp.1000genomes.ebi.ac.uk/vol1/ftp/data_collections/HGSVC2/working/20230515_Shi_hic_files/. Our *TRUHiC* method, along with the competing methods, the pre-trained *TRUHiC-LCL* model for chromosome 22, and all original code for the statistical analysis and pipeline implementation, have been deposited on GitHub https://github.com/shilab/TRUHiC and are publicly available as of the date of publication.

## Acknowledgment

This research includes calculations carried out on HPC resources supported in part by the National Science Foundation through major research instrumentation grant number 1625061 and by the US Army Research Laboratory under contract number W911NF-16-2-0189. We thank Rohan Alibutud for his feedback on the project and diligent proofreading.

## Human Genome Structural Variation Consortium (HGSVC)

The members of the Human Genome Structural Variation Consortium (HGSVC) are Hufsah Ashraf, Peter A. Audano, Olanrewaju Austine-Orimoloye, Parithi Balachandran, Anna O. Basile, Christine R. Beck, Marc Jan Bonder, Marta Byrska-Bishop, Mark J.P. Chaisson, Zechen Chong, André Corvelo, Jonathan Crabtree, Scott E. Devine, Peter Ebert, Jana Ebler, Evan E. Eichler (Co-Chair), Mark B. Gerstein, Bida Gu, Lisbeth A Guethlein, Pille Hallast, William T. Harvey, Patrick Hasenfeld, Alex R. Hastie, Mir Henglin, Kendra Hoekzema, PingHsun Hsieh, Sarah Hunt, Matthew Jensen, Miriam K. Konkel, Jan O. Korbel (Co-Chair), Jennifer Kordosky, Youngjun Kwon, Peter M. Lansdorp, Charles Lee (Co-Chair), Wan-Ping Lee, Alexandra P. Lewis, Chong Li, Jiadong Lin, Mark Loftus, Glennis A. Logsdon, Tobias Marschall (Co-Chair), Gianni V. Martino, Ryan E. Mills, Yulia Mostovoy, Mohammad Erfan Mowlaei, Katherine M. Munson, Giuseppe Narzisi, Andy Pang, David Porubsky, Timofey Prodanov, Keon Rabbani, Tobias Rausch, Xinghua Shi, Yuwei Song, Arda Söylev, Likhitha Surapaneni, Michael E. Talkowski, Vasiliki Tsapalou, Feyza Yilmaz, DongAhn Yoo, Xuefang Zhao, Weichen Zhou, and Michael C. Zody.

## HGSVC Functional Analysis Working Group

The members of the Human Genome Structural Variation Consortium (HGSVC) functional analysis working group are Anna O. Basile, Christine R. Beck, Marta Byrska-Bishop, Marc Jan Bonder, Mark J.P. Chaisson, Ken Chen, Evan E. Eichler, Matthew Jensen, Yunzhe Jiang, Kwondo Kim, Miriam K. Konkel, Jan O. Korbel (Co-Chair), Charles Lee, Chong Li, Jiaqi Li, Yang I. Li, Qingnan Liang, Glennis A. Logsdon, Tobias Marschall, Gianni V. Martino, Ryan E. Mills, Nicholas Moskwa, Yulia Mostovoy, Mark Gerstein, Lingbin Ni, Pille Hallast, Wolfram Höps, Daniel Ben-Isvy, Carolyn Paisie, Bernardo Rodriguez-Martin, Xinghua Shi (Co-Chair), Oliver Stegle, Sabriya Syed, Michael E. Talkowski, Yukun Tan, Alex Yenkin, DongAhn Yoo, Xuefang Zhao, Weichen Zhou, and Michael C. Zody.

